# Serum alpha-mannosidase as an additional barrier to eliciting oligomannose-specific HIV-1-neutralizing antibodies

**DOI:** 10.1101/2020.02.24.962233

**Authors:** Jean-François Bruxelle, Tess Kirilenko, Quratulain Qureshi, Naiomi Lu, Nino Trattnig, Paul Kosma, Ralph Pantophlet

## Abstract

Oligomannose-type glycans on HIV-1 gp120 form a patch that is targeted by several broadly neutralizing antibodies (bnAbs) and that therefore is of interest to vaccine design. However, attempts to elicit similar oligomannose-specific bnAbs by immunizing with oligomannosidic glycoconjugates have only been modestly successful so far. A common assumption is that eliciting oligomannose-specific bnAbs is hindered by B cell tolerance, resulting from the presented oligomannosides being sensed as self molecules. Here, we present data, along with existing scientific evidence, supporting an additional, or perhaps alternate, explanation: serum mannosidase trimming of the presented oligomannosides *in vivo*. Mannosidase trimming lessens the likelihood of eliciting antibodies with capacity to bind full-sized oligomannose, which typifies the binding mode of existing bnAbs to the oligomannose patch. The rapidity of the observed trimming suggests the need for immunization strategies and/or synthetic glycosides that readily avoid or resist mannosidase trimming upon immunization and can overcome possible tolerance restrictions.

## INTRODUCTION

Development of an effective HIV vaccine is still a high priority^1,2^. One of the central challenges remains the creation of a formulation that elicits broadly neutralizing antibodies (bnAbs) to the HIV envelope spike (Env). Six sites on Env are now recognized as vulnerable to bnAbs^3^, thus constituting vaccine targets. One of these sites is a conserved patch of oligomannose-type glycans on gp120. However, attempts to elicit bnAbs to this patch by immunization have not been fruitful.

The prevailing hypothesis is that tolerance mechanisms hinder the elicitation of oligomannose-specific bnAbs. Although the occurrence of oligomannose-type glycans is rare on healthy human tissue and cells^4^, a few human plasma glycoproteins do seem to sparsely express oligomannose-type glycans under normal physiological conditions^4^. The occurrence of these oligomannose-type glycans, even though not abundant, may be sufficient to limit the frequency of naïve B cells with receptors capable of binding oligomannose or render such ‘self-reactive’ B cells anergic^5^. Indeed, tolerance mechanisms are known to limit the repertoire frequency of naïve B cells with capacity to bind host glycan structures^6^. The identification of B cells with autoreactive signatures in HIV-infected individuals^7–9^, including those from whom oligomannose-specific bnAbs have been recovered^9^, has strengthened the notion that eliciting bnAbs, at least those to nominal self-glycans, may require tolerance restrictions to be overcome in some manner.

Most attempts to elicit oligomannose-specific bnAbs have involved the use of glycoconjugates with dense oligomannosyl clusters^10–17^. Dense clusters of oligomannose-type glycans are atypical of mammalian host protein glycosylation and thus were once thought suitable for defeating tolerance^18^. However, antibodies elicited by such approaches have been nearly invariably specific for oligomannoside substructures, rather than full-sized oligomannose as occurs on Env. These outcomes are interpreted commonly as a remaining manifestation of tolerance or conformational variances between natural oligomannose on Env and synthetic oligomannoside clusters^18,19^.

One possible alternate explanation for the prevalence of antibodies that are specific for oligomannose substructures—rather than full-sized oligomannose—could be that the elicited antibodies reflect a heretofore unappreciated occurrence *in vivo*: enzymatic trimming of the administered glycosides. This notion was raised in a recent report^20^ describing an attempt to elicit 2G12-like bnAbs in rabbits using multivalently displayed glycopeptide immunogens. The report showed that the resulting antibodies were largely specific for the proximal glycan core rather than the distally located tips of the target glycans^20^. It was suggested that this outcome might be the result of the activity of a mannosidase in (rabbit) serum that trims full-sized oligomannose on administered glycoproteins^21–23^. Serum mannosidase trimming has been reported to occur fairly rapidly (*t*_½_ ~5-6 h)^21,22^; if so, then that would readily decrease the availability of full-sized oligomannosides for recognition by B cells at priming and subsequent antibody affinity maturation in germinal centers^24^. The significance of such enzymatic trimming is substantial given that bnAbs to the oligomannose patch on HIV gp120 are specific for full-sized oligomannose^9,25–28^ and considering that glycans are a major component of the epitope of many other bnAbs.

Here, we report on our own investigation of serum mannosidase as an additional potential hurdle to eliciting oligomannose-specific bnAbs. First, we show that bnAb PGT128, an example of the newer generation of bnAbs to the oligomannose patch, binds substantially worse to a microtiter plate-bound glycoconjugate that has been incubated in situ with mammalian sera, with sera from mice and humans notably causing the greatest reduction. Antibody binding is restored when kifunensine, a highly specific alpha-mannosidase inhibitor^29,30^, or EDTA is added to the serum, demonstrating that the loss of antibody binding is due to Ca^2+^-dependent alpha-1,2-specific mannosidase activity. Secondly, and perhaps more significantly, we show that early antibodies produced in animals immunized with a CRM_197_-conjugated glycomimetic of oligomannose^31^ bind better to serum-treated glycoconjugate (i.e., enzyme-trimmed) than to buffer-treated glycoconjugate, suggesting that glycoside trimming indeed occurs *in vivo*. In addition to having obvious implications for the design of glycoconjugates for eliciting oligomannose-specific bnAbs, we show that our findings are also of relevance to efforts employing recombinant envelope glycoproteins.

## RESULTS

### Bacterially derived glycomimetic of mammalian oligomannose conjugated to protein carrier CRM_197_ retains favorable antigenicity

We have reported previously on a synthetic antigenic mimic of mammalian oligomannose^31^, which was designed based on the lipooligosaccharide backbone of an *Rhizobium radiobacter* strain that closely resembles the D1 arm of oligomannose^32,33^ and then synthetically extended to create an analog of the D3 arm of mammalian oligomannose. For practical purposes, the mimetic was conjugated initially to BSA^31^. We subsequently conjugated it to CRM_197_, a more clinically apt carrier protein ^34^ that stimulates robust T-follicular helper (Tfh) responses^35–38^. As before^31^, the glycomimetic, equipped at the reducing end with an amine linker, was conjugated to lysine residues present on the protein carrier, via an isothiocyanate intermediate. MALDI-TOF analyses showed that, depending on the batch, 3.5 – 6.5 glycosides could be conjugated per CRM_197_ molecule (Supplementary Fig. S1). We evaluated the ability of bnAb PGT128 and three related members of the PGT128/130 bnAb family to bind this new CRM_197_ glycoconjugate, which was named NIT211. As shown in Fig. 1, all four bnAbs bind NIT211 at least as good as NIT82B, the initial BSA conjugate^31^.

In summary, our results show that the CRM_197_-conjugate NIT211 reported here presents a reasonable mimic of oligomannose as occurs on HIV Env, as evidenced by reasonably strong binding of bnAb PGT128 and related antibodies.

**Figure 1.**
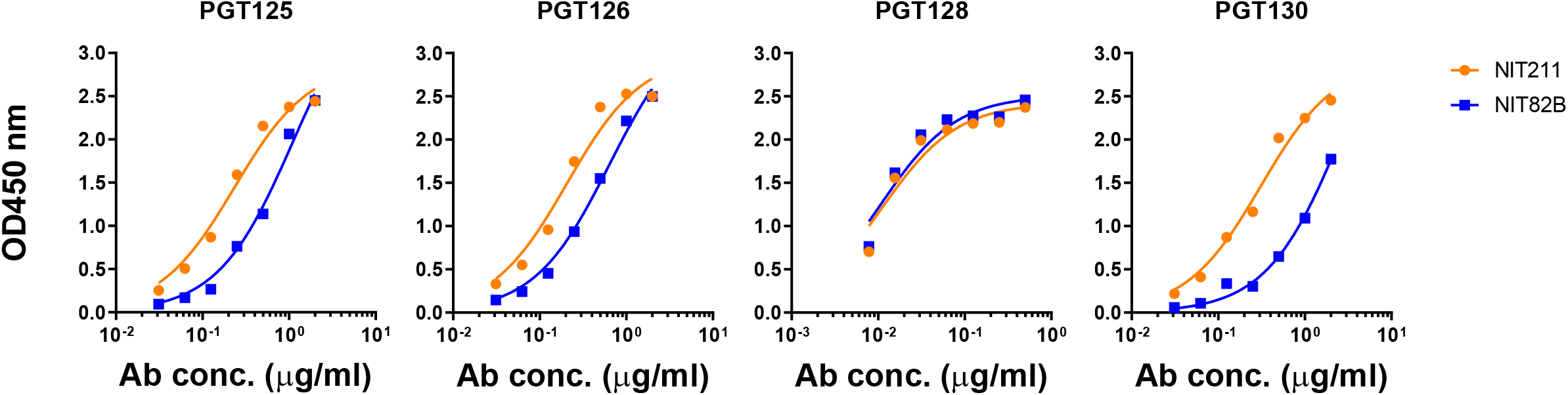
PGT128 and related bnAbs bind the CRM_197_-conjugated mimetic (NIT211) with similar or greater avidity as the BSA-conjugated version (NIT82B). The NIT211 derivative (NIT211_3) used here is loaded at 3.5 glycosides per CRM_197_. NIT82B is loaded at 4.4 glycosides per BSA. The conjugates (63-73 kDa) were coated as solidphase antigen onto microtiter-plate wells at 5 μg/ml and assayed for recognition by PGT125, 126, 128, and 130. All antibodies were tested as IgGs.

### Serum mannosidase trims the CRM_197_-conjugated glycoside *in vitro*

In the recent report suggesting that serum mannosidases might be impeding the elicitation of oligomannose-specific bnAbs^20^, the Man_9_-peptide conjugate designed in the study was retrieved for mass spectrometric analysis after incubation in rabbit serum. Results showed that >90% of the Man_9_ was trimmed to Man_6_ within 24 h. In line with the results of that report, we found here using an ELISA format that PGT128 binding to both the BSA-conjugated (NIT82B) and the CRM_197_-conjugate glycoside (NIT211) is substantially diminished after a 24 h-incubation in situ with different mammalian sera relative to buffer control (Fig. 2). Notably, the reduction in antibody binding was particularly pronounced following incubation in human and mouse sera compared to rat or rabbit sera. Adding EDTA or kifunensine to these sera restored PGT128 binding (Fig. 2), consistent with the enzymatic activity of a Ca^2+^-dependent alpha-1,2-specific mannosidase^21^.

**Figure 2.**
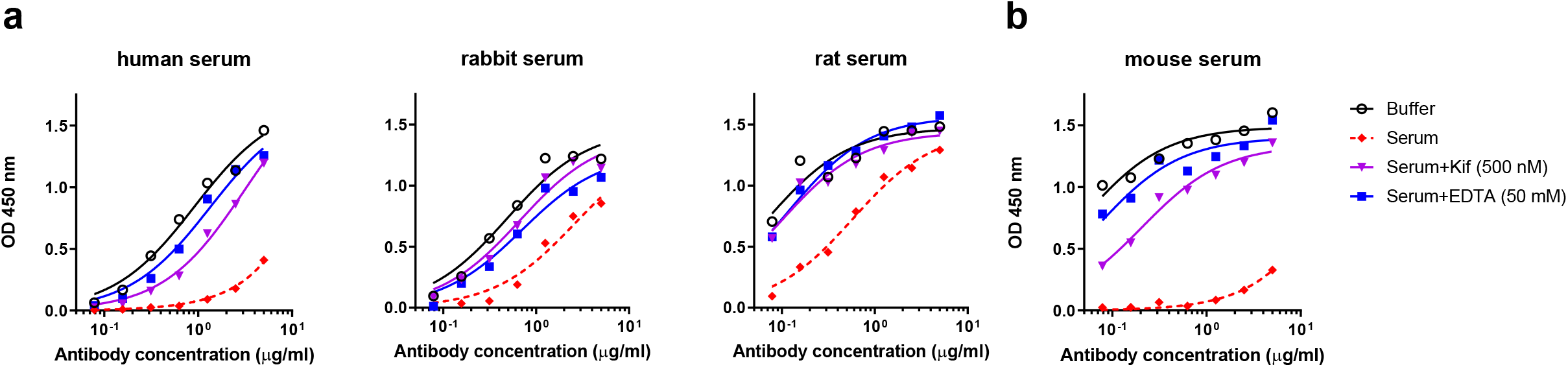
Serum mannosidase trims protein-conjugated oligomannose mimetic *in vitro*. Shown is PGT128 binding to the BSA-conjugated (**a**) or the CRM_197_-conjugated glycoside (**b**) (5 μg/ml) after in situ overnight incubation of glycoconjugate-coated ELISA plates with buffer, mammalian serum, serum supplemented with kifunensine (Kif) or serum supplemented with EDTA. All experiments for a given serum were performed on a single assay plate to avoid potential plate-to-plate variability. Shown are representative results from two independent experiments.

We performed assays to ascertain that reduced PGT128 binding was due to enzymatic activity in serum and not an artifact of our set up. First, we confirmed that the restoration of antibody binding with both kifunensine and EDTA was titratable (Supplementary Fig. S2); PGT128 binding increased with increasing inhibitor concentration (i.e., increased inhibition of mannosidase directly (kifunensine) or sequestration of the ionic cofactor Ca^2+^ (EDTA)). Second, the reduction in PGT128 binding following incubation of the glycoconjugate with serum could also be lessened in the presence of deoxymannojirimycin (DMJ) (Supplementary Fig. S2), another alpha-mannosidase inhibitor. These results further strengthen the case for mannosidase activity in serum leading to the observed reduction in PGT128 binding. The inhibitory effect of DMJ was less pronounced than with kifunensine, which is consistent with the reported weaker potency of DMJ against alpha-1,2-mannosidases compared to kifunensine^30^.

### Serum mannosidase may trim oligomannosidic glycoconjugates *in vivo*

Having confirmed mannosidase activity in serum, we sought also to determine whether serum mannosidase trimming is relevant *in vivo*. To do so, we used sera from Trianni mice that were immunized three times (days 0, 21 and 42) with adjuvanted NIT211, the CRM_197_ conjugate of our oligomannose mimetic. Trianni mice express a complete human antibody repertoire, thus enabling an approximation of potential antibody responses in people. We assayed sera collected 7 days after the second booster immunization (day 49) for binding to NIT82B, the BSA conjugated version of the mimetic, which had been incubated beforehand (overnight) in situ with buffer or human serum. As shown (Fig. 3 and Supplementary Fig. S3), sera that exhibited measurable binding to the BSA-glycoconjugate tended to bind better to the serum-treated conjugate than to the buffer-treated conjugate. These results are suggestive of serum mannosidase trimming, at least to some extent, of (synthetic) glycosides conjugates *in vivo* upon immunization.

**Figure 3.**
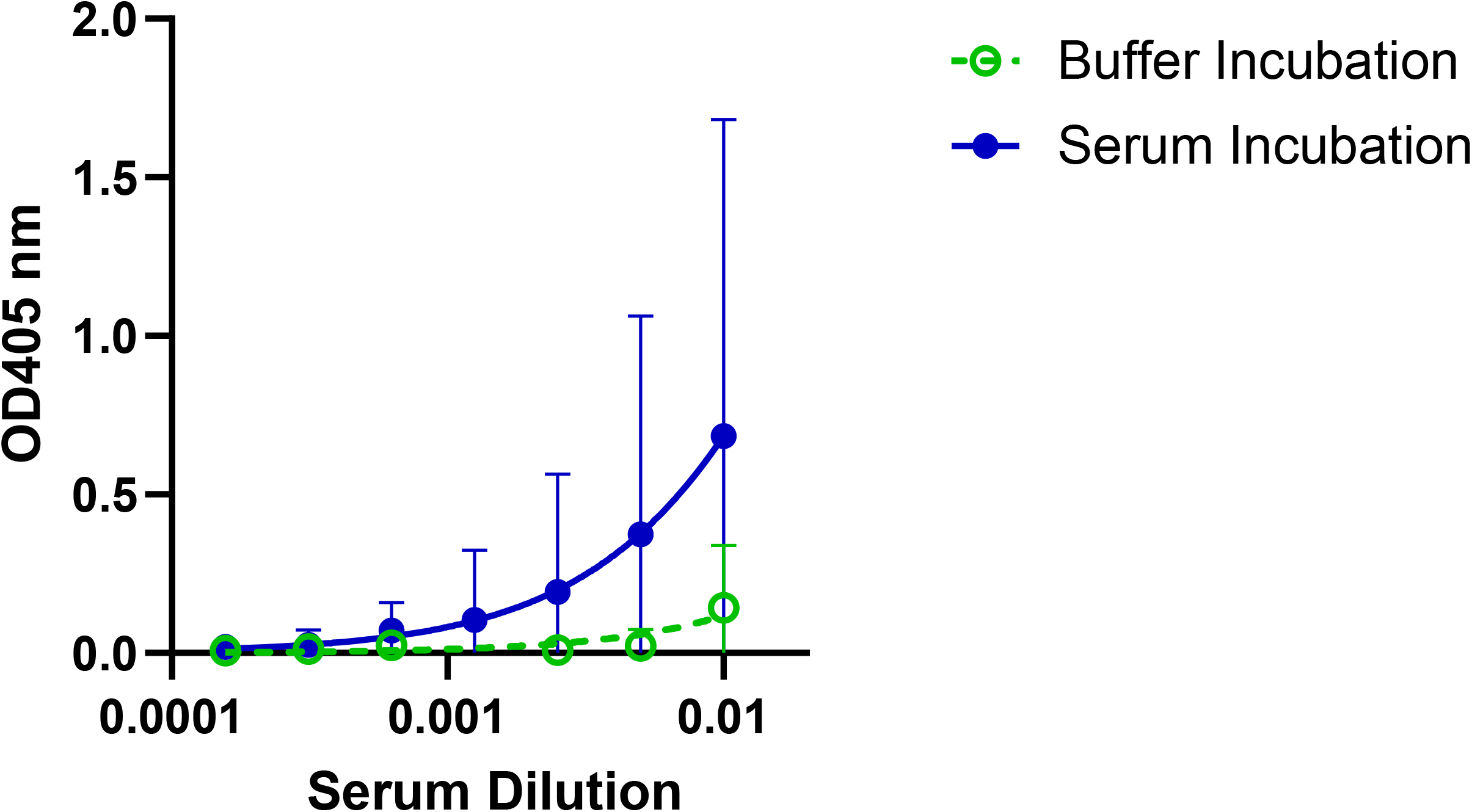
Sera from Trianni mice immunized with the adjuvanted CRM_197_ conjugate of our oligomannose mimetic bind preferentially to the serum mannosidase-trimmed glycoconjugate. Sera collected on day 49 from Trianni mice (n=5) that were immunized three times (days 0, 21, 42) with GLA-SE adjuvanted CRM_197_-oligomannoside conjugate NIT211 (pre-made mixture of NIT211_4 (5.9 ligands/CRM_197_) and NIT211_5 (6.5 ligands/CRM_197_), were assayed for binding to a BSA glycoconjugate (NIT82B)-coated ELISA plate after overnight (24 h) incubation of the glycoconjugate-coated wells with buffer or with human serum. Graphs depict geometric mean values for the five serum samples, each assayed in duplicate, with error bars denoting the standard deviation from the mean.

We did observe that not all serum samples bound the buffer-treated BSA conjugate at the serum dilutions assayed here (starting at 1:100) (Supplementary Fig. S3). These sera were nevertheless included to determine whether binding to the serum-treated BSA-conjugate (i.e., trimmed glycoside) would be observed – which we however did not. The absence of glycoside-binding antibodies in these sera does not appear to be the result of poor immunogenicity of the administered glycoconjugate per se, given the robust antibody titers to the carrier protein (CRM_197_) observed with all animals (Supplementary Fig. S3).

### Serum alpha-mannosidase does not trim oligomannose-type glycans on recombinant envelope glycoprotein trimers

The results above led us to consider that serum mannosidase might also act on recombinant envelope glycoprotein trimers. So-called SOSIP trimers in particular have gained substantial interest in the last several years as vaccine immunogens^39^. SOSIP trimer glycosylation does not exactly match glycosylation on the native virus spike; SOSIP trimers tend to possess a somewhat higher proportion of oligomannose-type glycans^40,41^, possibly because of quaternary structure differences between SOSIP trimers and the native spike^42,43^. Nevertheless, given the strong interest in using SOSIP and SOSIP-like trimers to elicit bnAbs and progress in the development of other recombinant HIV envelope glycoprotein trimers, determining how exposure to serum mannosidase might affect the presented glycans seems warranted. We found however that there was no significant difference in PGT128 binding to SOSIP trimers ZM197M and C.ZA97^44,45^ that were incubated overnight in situ with human serum compared to the buffer-treated controls (Fig. 4). No change in antibody binding was observed upon the addition of kifunensine. Thus, these results show that, in contrast to glycoconjugates, serum alpha-mannosidase does not trim oligomannose presented on recombinant envelope glycoproteins that closely resemble the native spike. Serum mannosidase therefore presumably also does not trim clustered oligomannose on the native glycoprotein spike.

**Figure 4.**
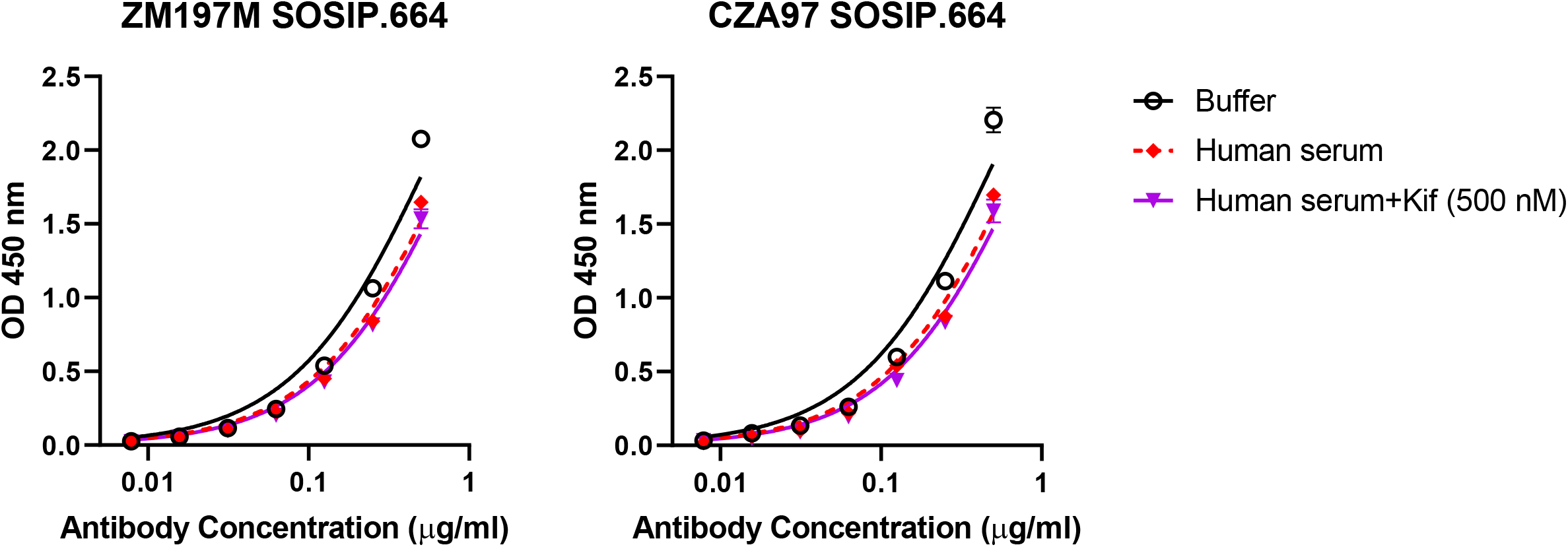
SOSIP trimers are refractory to serum mannosidase trimming. Binding of a murine (IgG2a) version of antibody PGT128 to HIS-tagged SOSIP trimers ZM197M and C.ZA97 absorbed onto Ni^2+^-coated plates (5 μg/ml) was determined following in situ overnight incubation with buffer or human serum. Results are from a single experiment.

## DISCUSSION

Despite substantial effort, attempts to elicit antibodies to the oligomannose patch on HIV have not truly made much progress to date. Although B cell tolerance is often thought to be the major impediment –and may still be relevant—other factors have received relatively little attention to date. Following from a recent study^20^ in which it was suggested that serum alpha-mannosidase trimming may (also) be an impediment to these efforts, in this report we provide further evidence in support of this thesis. Specifically, our results suggest that serum mannosidase trimming may occur readily following immunization with oligomannosidic glycoconjugates (Fig. 3).

In contrast to our observations with the conjugates, *in vitro* assays with SOSIP trimers showed no trimming of glycans comprising the oligomannose patch, demonstrating that these oligomannose-type glycans, unlike those presented on glycoconjugates, are protected from mannosidase trimming, presumably because of their more dense packing on Env. Our results along with those of Nguyen et al^20^ allow for an explanation of why immunizing rabbits with recombinant yeast expressing an abundance of oligomannose^16,46,47^ or prolonged repetitive immunization of macaques with a recombinant gp140 trimer imparted with an intact oligomannose patch ^48^ might yield mannose-specific antibodies with at least some affinity for full-sized oligomannose, whereas immunizing with oligomannose or oligomannose-like glycans displayed on virus-like particles^11,49^, oligomannosidic glycoconjugates^10,12,13,15,17^ or glycopeptides presenting oligomannose^20,28^ has resulted mostly in the elicitation of antibodies that are generally less capable of binding oligomannose.

The physiological role of serum alpha-mannosidase is not entirely clear. One seemingly plausible suggestion^21^ is that it may be responsible for demannosylating serum glycoproteins or plasma membrane glycoproteins, which would explain why oligomannose-type glycans are relatively rare on mammalian cells and tissue^4^ – lest they perhaps inadvertently trigger complement activation via the mannose-binding lectin pathway. Although it is not yet clear where serum mannosidase acts upon an administered antigen *in vivo*, one seemingly plausible notion is that it is present in transudated plasma entering lymph nodes where it trims antigen poised for immune recognition. Unknown also is whether serum mannosidase levels vary in people; although little variation in serum mannosidase levels has been reported in British people^50^, it is unknown whether the same is true in other populations.

How can we reconcile the development of oligomannose-specific bnAbs with alpha-mannosidase activity in serum? Clearly, the results above with the SOSIP trimers (Fig. 4) indicate that serum mannosidase does not easily trim clustered oligomannose-type glycans on Env, meaning that these glycans would be available for recognition by potential B cells during infection. Indeed, experimental evidence suggests that a sustained presence of envelope glycoprotein with the ‘proper’ glycosylation over several weeks (or months) of infection might trigger the ‘right’ B cells. ^51,52^. Of course, we do not know which form(s) of the envelope glycoprotein trigger(s) the development of oligomannose-specific bnAbs during infection^53^ or whether the triggered B cells are nominally autoreactive or anergic^54^. Further investigations, perhaps such as done for antibodies to the membrane-proximal region of gp41^55^, may help to answer these questions.

The results presented here might seem to favor the use of SOSIP-based trimers rather than, for example, neoglycoconjugates to elicit bnAbs to (oligomannose-type) glycans on HIV. For example, one study reported on the elicitation of glycan-dependent antibodies resembling precursors of human bnAbs to the oligomannose patch in non-transgenic animals using a ‘designer’ SOSIP-based trimer^56^. Despite potential advantages of SOSIP-based immunogens, one probable disadvantage is glycosylation heterogeneity, which has been experimentally shown to be greater at several *N*-glycosylation sites compared to native Env^40^. Synthetic oligomannosides offer an attractive alternative in this regard, enabling production of structurally defined immunogens. Another distinct advantage, and the one on which our glycomimetic approach is largely based, is that synthetic oligomannosides can be designed to overcome potential hindrance in B cell activation; autoreactive B cells can be ‘awakened’ when they are suddenly exposed to cross-reactive antigens in a milieu with the proper immunostimulatory signals^54^.

In conclusion, we confirm and expand here on recent research showing the heretofore unappreciated occurrence of a soluble alpha-1,2-mannosidase in mammalian serum that may be responsible, at least in part, for blunting the ability of past glycoconjugate approaches to elicit oligomannose-specific responses. Our results suggest that the mannosidase activity is particularly strong in human and mouse sera; it seems less pronounced in rat and rabbit sera, perhaps explaining why we were previously able to elicit modest levels of oligomannose-specific antibodies in transgenic rats^31^. Identifying one or more approaches that can best overcome this mannosidase activity is an obvious next step. Given the rapidity with which trimming occurs (>90% trimming of Man_9_ within 24 h^20–22^), it is clear that any forthcoming approach will need to be able to prevent or at least minimize mannosidase activity immediately upon administration of the immunogen to allow enough time for B cell activation at priming, which for T-dependent primary antibody responses is typically delayed (~2-3 days postimmunization) due to the need for sufficient recruitment of T-cell help for B cell activation to occur^57,58^. Therefore, parallel to probing immunization strategies to limit mannosidase trimming, we intend to explore the design of mannosidase-resistant glycosides to provide the highest possible resistance to this unforeseen hindrance. Given that antibodies to the oligomannose patch on HIV-1 are major contributors to neutralization breadth and potency in many HIV-infected individuals^59–61^, we feel that these efforts are worthwhile as part of the broader endeavor of developing an effective HIV vaccine component to elicit bnAbs.

## METHODS

### Glycoside synthesis

The soluble heptamannoside mimetic of oligomannose used for these studies, termed NIT68B, was synthesized as described previously^31^.

### General procedure for neoglycoconjugate preparation

The BSA conjugate of ligand NIT68B, with an average loading density of 4 glycosides per BSA molecule, was prepared as reported^31^. The CRM_197_ conjugates were synthesized as described recently^62^ but with minor modifications. The amine ligand NIT68B (3.1 mg, 2.6 μmol) in 0.1 M NaHCO_3_ (1 ml) was vigorously stirred with a solution of thiophosgene (3.9 μl, 51 μmol) in chloroform (1 ml) for 4 h at RT. The organic phase was removed and extracted 4 times with CHCl_3_. Traces of organic solvent were removed from the aqueous phase by bubbling a stream of air through the solution. Then a solution of CRM_197_ (1.0 mg, 17.1 nmol) in 0.3 M NaCl/0.1 M NaHCO_3_ (0.5 ml) was added and stirring was continued for 72 h at RT. The solution was passed through an Amicon Spin filter (10 kDa) to remove unreacted ligand and washed with PBS buffer to give a solution of the neoglycoconjugate NIT211_3 (0.7 ml). The protein concentration (0.36 mg/ml) was determined using extinction data obtained with a Nanodrop instrument. Conjugate NIT211_4 was prepared using 4.0 mg (3.3 μmol) NIT68B in 0.1 M NaHCO_3_ (0.5 ml), thiophosgene (10.1 μl, 0.132 mmol) and CRM_197_ (2.0 mg, 34.2 nmol) in 0.3 M NaCl/0.1 M NaHCO_3_ (0.7 ml) and was obtained as a solution of the neoglycoconjugate (1 ml; 1.85 mg/ml). Conjugate NIT211_5 was prepared using 12.0 mg (9.9 μmol) NIT68B in 0.1 M NaHCO_3_ (1.0 ml), thiophosgene (30.4 μl, 0.397 mmol) and CRM_197_ (6.0 mg, 102.7 nmol) in 0.3 M NaCl/0.1 M NaHCO_3_ (1.0 ml) and was obtained as a solution of the neoglycoconjugate (1.5 ml; 3.44 mg/ml). For all conjugates, the amount of conjugated ligand per carrier molecule was determined by MALDI-TOF mass spectrometry. MALDI data showed an average loading density of 4.4 glycosides per BSA molecule^31^ and 3.5-6.5 glycosides per CRM_197_ molecule (Supplementary Fig. S1).

### Antibody expression and purification

Antibodies PGT125, 126, 128 and 130 were expressed from FreeStyle 293F cells grown in FreeStyle media supplemented with 2% FBS following transfection at a 1:1 ratio of expression plasmids pFUSEss-CLIg-hL2 and pFUSEss-CHIg-hG1 encoding the light and heavy chain, respectively, of each corresponding antibody. The variable light and heavy chains of each antibody were codon-optimized (ATUM) for expression in 293 cells. A murine version of PGT128 was expressed also, using expression plasmids pFUSE2ss-CLIg-mL1 and pFUSEss-CHIg-mG2a encoding the light and heavy chain, respectively, of the antibody. For transfections, plasmids were mixed with either 293fectin, 293-free or ExpiFectamine 293 diluted in Opti-MEM I media; all three agents are optimized for transfection of HEK293-type cells grown in suspension culture. Cell culture supernatants were harvested 6 days after transfection, filtered, concentrated and antibody then purified on individual protein A (human IgG) or protein G (mouse IgG) spin columns as per the recommended protocols. Sample purity was verified by SDS-PAGE and antibody concentration determined on a Nanovue spectrophotometer. Prior to using the murine version of PGT128, we confirmed that it binds antigen just as well as the human version of the antibody (Supplementary Fig. S4).

### Trianni mouse immunizations

Trianni mice (n=5, mix of male (3) and female (2) animals; 6-8 weeks of age at the start of immunization) were immunized under contract at Antibody Biosolutions (Sunnyvale, CA). For immunization, a ~1:3 mixture consisting of NIT211 derivatives NIT211_4 (5.9 ligands/CRM_197_) and NIT211_5 (6.5 ligands/CRM_197_) was used. The mixed NIT211 conjugate (30 μg, corresponding to ~3 μg of conjugated glycoside) was mixed with an equal amount of glucopyranosyl lipid A in a stable emulsion (GLA-SE), and incubated for 1 h at room temperature before being injected subcutaneously into each animal. The experimental protocol (protocol. no. 1242HS-17) for using mice in this study was approved by the University Animal Care Committee (UACC) of Simon Fraser University and followed relevant guidelines and regulation of animal care. A small bleed was collected from all animals immediately prior to immunization and at 7 days after the boosters at days 21 and 42. The collected blood was left to clot so that serum could be recovered. Serum samples were stored at −20 °C. Once thawed, they were kept at 4 °C.

### ELISA

ELISAs were performed as described previously^31^. For the serum mannosidase trimming experiments, buffer, serum (neat) or serum plus inhibitor were added after the blocking step and incubated overnight (24 h) in a humidified incubator at 37 °C. The next day, the plate was washed and the assay continued as per standard with the primary antibody. For the ELISAs with glycoconjugates, polystyrene microtiter plates were used whereas nickel-coated plates were used for SOSIP trimer ELISAs. Tetramethylbenzidine (TMB) was used as substrate; the reaction was stopped with H2SO4 (2 M) and read immediately at 450 nm.

### Software/Data analysis

All ELISA data was graphed with Graphpad Prism.

## ACKNOWLEDGMENTS

The research described in this report was supported by funding from the Austrian Science Fund (P26919-N28 to P.K.), the Canadian Institutes of Health Research (IBC-150408 to R.P.), and the National Institutes of Health (R01 AI134299 to R.P.). R.P. was supported also by a salary award from the Michael Smith Foundation for Health Research (no. 5268). T.K. was supported by a GlycoNet summer award for undergraduate students. We thank Darrick Carter and Shari Maxwell (Infectious Disease Research Institute) for access to GLA-SE adjuvant, Matthias Wabl and Maria Wabl (Trianni Inc) for access to Trianni transgenic mice, and Billy Nguyen and Glen Lin (Antibody Solutions) for coordinating the immunizations. We are particularly grateful to John Moore, Albert Cupo and Victor Cruz Portillo (Weill Cornell Medical College) for generously providing the C.ZA97 SOSIP trimer and to Rogier Sanders, Ron Derking and Tom Bijl (Amsterdam UMC) for the ZM197M SOSIP trimer. We also thank Asa Lau and Kurtis Ng for technical assistance.

## AUTHOR CONTRIBUTIONS

Conceptualization – R.P.; Formal analysis – J.-F., N.T., P.K., R.P.; Funding acquisition – P.K., R.P.; Investigation – J.-F., T.K., Q.Q., N.L., N. T.; Supervision – J.-F., P.K., R.P.; Validation – P.K., R.P.; Writing – original draft – J.-F., R.P.; Writing – review & editing – J.-F., T.K., Q.Q., N.L., N.T., P.K., R.P.

## ADDITIONAL INFORMATION

No competing interests.

