## Supplemental Information for "Serum alpha-mannosidase as an additional barrier to eliciting oligomannose-specific HIV-1-neutralizing antibodies"

^1^Faculty of Health Sciences, Simon Fraser University, Burnaby, British Columbia V5A1S6, Canada; ^2^Department of Molecular Biology & Biochemistry, Simon Fraser University, Burnaby, British Columbia V5A1S6, Canada; ^3^Department of Chemistry, University of Natural Resources and Life Sciences, Vienna A-1190, Austria; ^4^Present address: AbCellera Biologics Inc., Vancouver, British Columbia, Canada; ^5^Present address: National Collaborating Centre for Infectious Diseases, Winnipeg, Manitoba, Canada; ^6^Present address: Department of Chemical Biology and Drug Discovery, Utrecht University, Utrecht, The Netherlands

**Figure S1.** MALDI-TOF spectra of NIT211 derivatives. *Top:* Compounds NIT211_3, NIT211_4 and NIT211_5 were synthesized in essence as described recently^1^, albeit with a few alterations as detailed here in the Methods section. The aminopropyl-equipped oligomannoside was first activated with thiophosgene to form a isothiocyanate intermediate and was then added to a solution of CRM_197_ in buffer to furnish the neoglycoconjugate. A higher dilution of reagents during conjugation resulted in a lower carbohydrate density of the corresponding glycoconjugates. *Bottom:* MALDI-TOF spectra of CRM_197_, NIT211_3, NIT211_4 and NIT211_5 (top to bottom, respectively). The MALDI-TOF mass spectrometric analysis revealed a ligand:CRM_197_ ratio of 3.5:1 (NIT211_3), 5.9:1 (NIT211_4) and 6.5:1 (NIT211_5).

**Figure S2.** Inhibition of serum mannosidase trimming by kifunensine and EDTA *in vitro* is titratable and can be achieved with deoxymannojirimycin (DMJ), another alpha-mannosidase inhibitor. (**a**) Titratable inhibition of serum mannosidase trimming *in vitro*. Shown is PGT128 binding to the BSA-conjugated glycoside (at fixed concentration (5 µg/ml)) after in situ overnight incubation of glycoconjugate-coated ELISA plates with rat or mouse containing graded concentrations of kifunensine (left) or EDTA (right). The test concentrations for each inhibitor were performed on a single assay plate to avoid potential plate-to-plate variability. The data show that PGT128 binding increases with increasing inhibitor concentration. (**b**) Inhibition of serum mannosidase trimming by DMJ. Shown is the binding of serially titrated antibody PGT128 to the BSA-conjugated oligomannoside mimetic NIT82B after in situ overnight incubation of a glycoconjugate-coated (5 µg/ml) ELISA plate with buffer, rat serum, rat serum supplemented with the potent alpha-1,2-mannosidase inhibitor kifunensine (Kif) or rat serum supplemented with the alpha-1,2-mannosidase inhibitor DMJ.

**Figure S3.** Binding of sera from individual Trianni mice to the BSA-conjugated oligomannose mimetic NIT82B in comparison to serum binding to the CRM_197_ protein carrier. *Left:* Sera collected on day 49 from five Trianni mice immunized three times (days 0, 21, 42) with a GLA-SE adjuvanted formulation of our oligomannose mimetic conjugated to CRM_197_ were assayed for binding to a BSA-conjugate of the same glycoside (NIT82B) which had been coated onto ELISA plate wells overnight and subsequently incubated an additional night (24 h) with buffer or with human serum. Each serum was assayed in duplicate, with error bars denoting the standard deviation from the mean. Denoted above each graph is also the sex of each of the five animals. *Right:* ELISA of pre-immune sera (day 0; D0) and immune sera (day 49; D49) from the five individual Trianni mice to the CRM_197_ protein carrier, which was coated onto microtiter plate wells at 5 μg/ml.

**Figure S4.** Human and mouse versions of bnAb PGT128 bind the oligomannose mimetic NIT82B equivalently. Human IgG1 (blue curve) and mouse IgG2a (red curve) versions of bnAb PGT128 were assayed for binding to BSA glycoconjugate NIT82B coated as solid-phase antigen (5 μg/ml) onto ELISA plate wells.
