## Supplementary figures and images for "Serum alpha-mannosidase as an additional barrier to eliciting oligomannose-specific HIV-1-neutralizing antibodies"

### Supplemental Figure 1

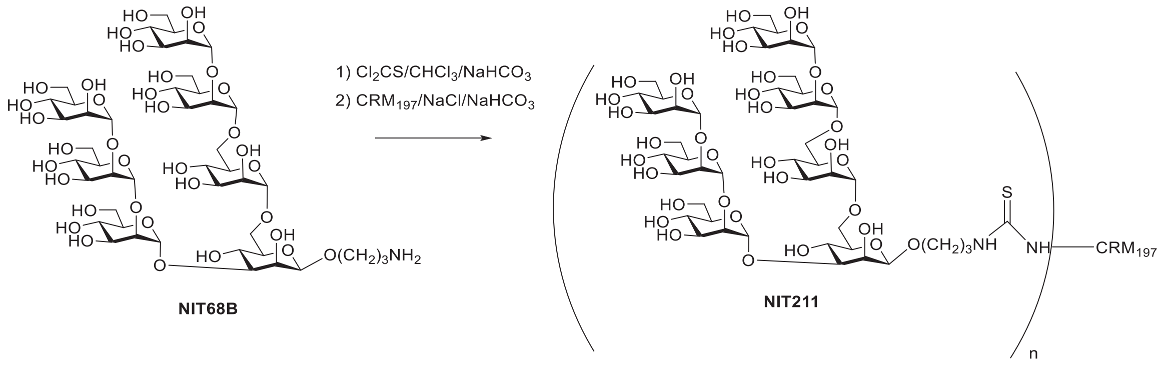

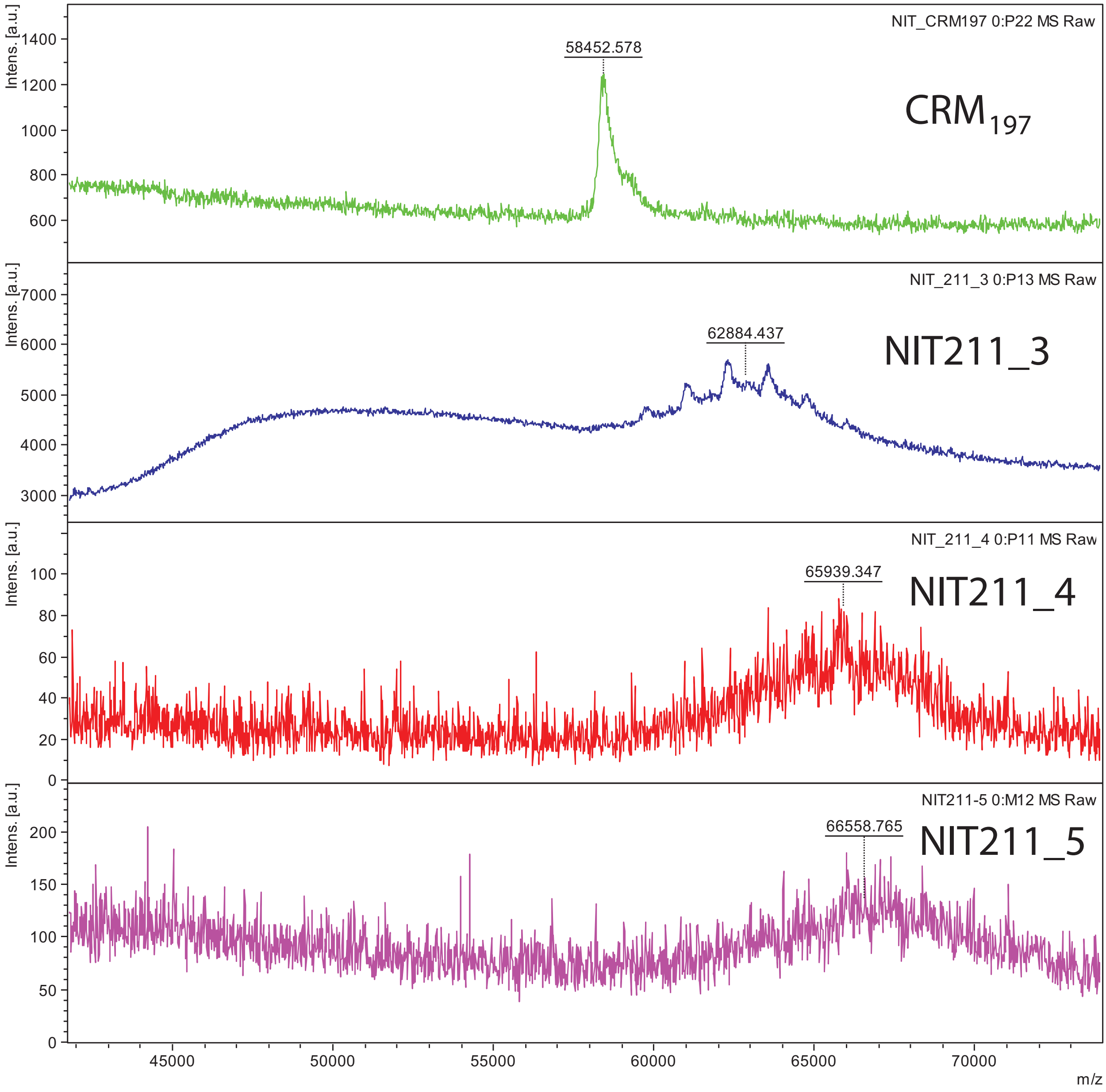

### Supplemental Figure 2

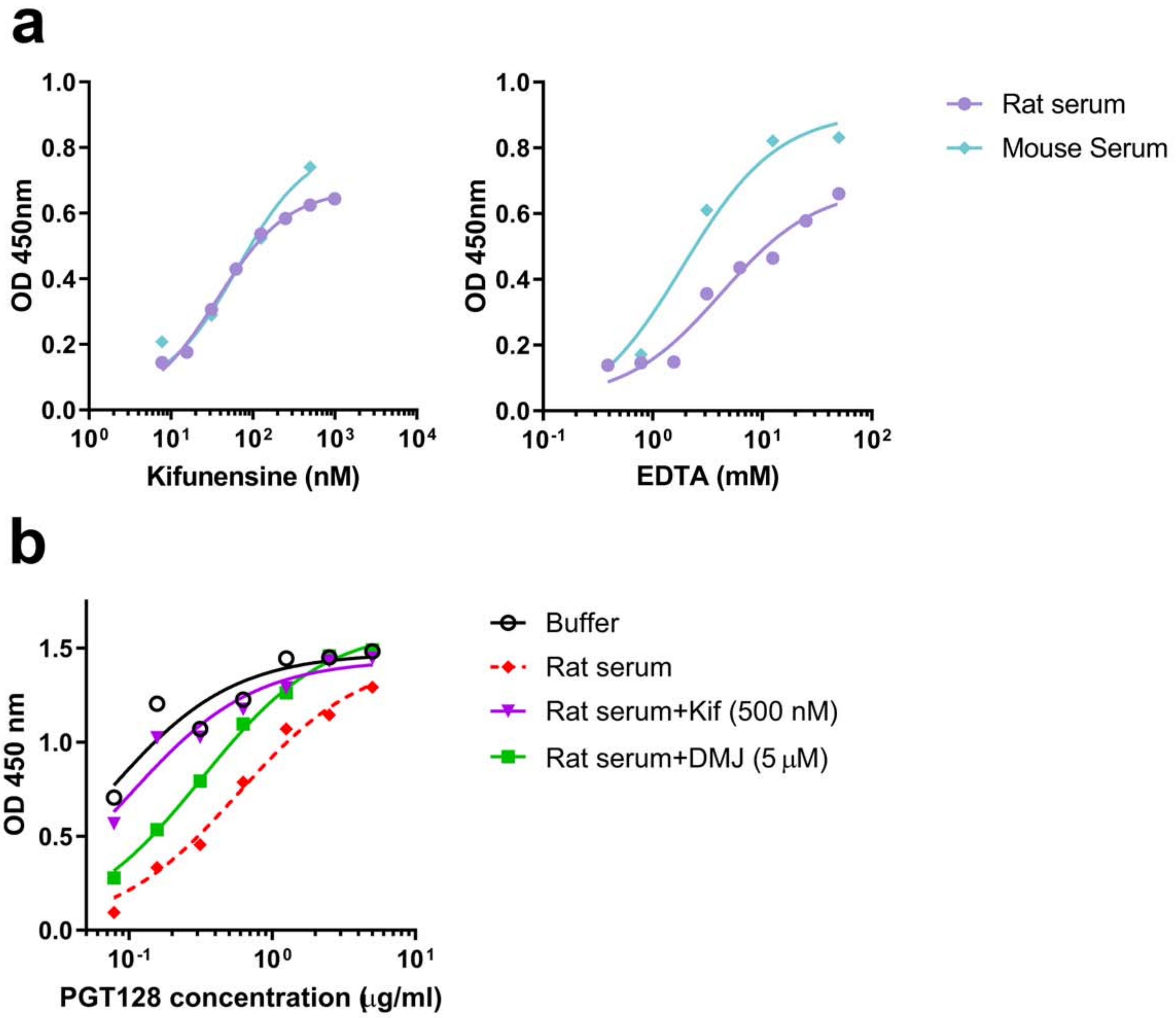

### Supplemental Figure 3

#### BSA-conjugate

#### $CRM_{197}$ protein carrier

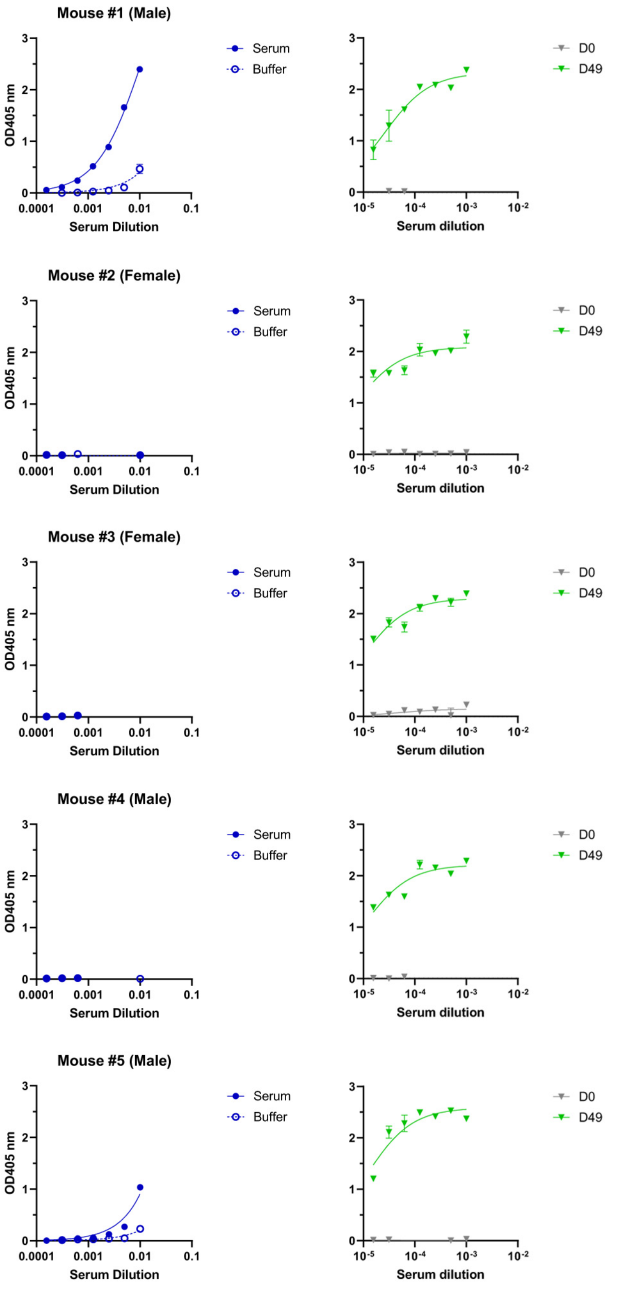

### Supplemental Figure 4

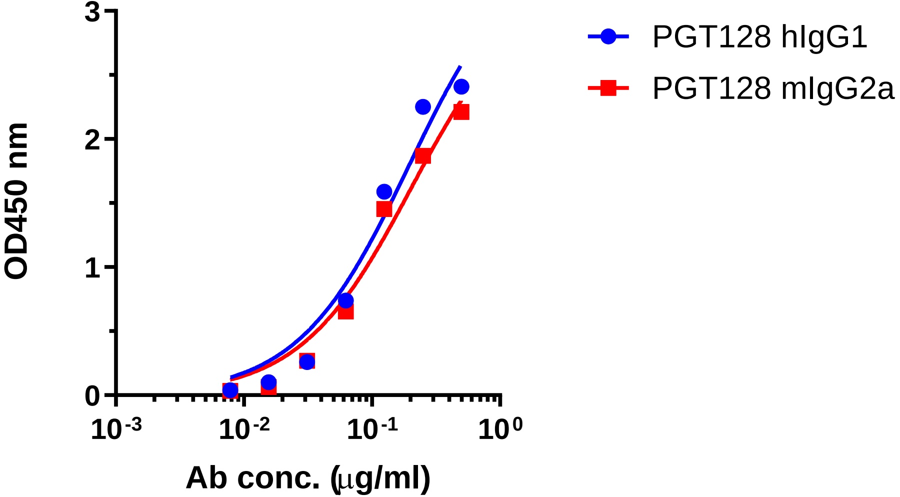
